# Characterisation of CASPR2 deficiency disorder - a syndrome involving autism, epilepsy and language impairment

**DOI:** 10.1101/034363

**Authors:** Pedro Rodenas-Cuadrado, Nicola Pietrafusa, Teresa Francavilla, Angela La Neve, Pasquale Striano, Sonja C. Vernes

**Author notes:** Corresponding author contact details: Sonja C. Vernes.

## Abstract

**Background:** Heterozygous mutations in *CNTNAP2* have been identified in patients with a range of complex phenotypes including intellectual disability, autism and schizophrenia. However heterozygous *CNTNAP2* mutations are also common in the normal population. Conversely, homozygous mutations are rare and have not been found in unaffected individuals.

**Case presentation:** We describe a consanguineous family carrying a deletion in *CNTNAP2* predicted to abolish function of its protein product, CASPR2. Affected family members show epilepsy, facial dysmorphisms, severe intellectual disability and impaired language. We compared these patients with previously reported individuals carrying homozygous mutations in *CNTNAP2* and identified a highly recognisable phenotype.

**Conclusions:** We propose that CASPR2 loss produces a syndrome involving early-onset refractory epilepsy, intellectual disability, language impairment and autistic features that can be recognized as CASPR2 deficiency disorder. Further screening for homozygous patients meeting these criteria, together with detailed phenotypic investigations will be crucial for understanding the contribution of *CNTNAP2* to normal and disrupted development.

## INTRODUCTION

The *CNTNAP2* gene (located on 7q35) is encoded by 24 exons, spanning 2.3 Mb, making it a large physical target for disruptive mutations. The majority of reported patients carry heterozygous disruptions of *CNTNAP2* and usually display some combination of core phenotypes including intellectual disability (ID), seizures, autistic features and impaired language [1]. However, the phenotype associated with heterozygous *CNTNAP2* mutation is complex as the severity of characteristics varies across patients, and in some cases additional phenotypes are observed (e.g. Tourette syndrome, schizophrenia). Furthermore, coding changes in a single copy of *CNTNAP2* have also been found in unaffected individuals, suggesting heterozygous mutations are not always fully penetrant.

*CNTNAP2* encodes the CASPR2 protein - a member of the neurexin superfamily of proteins that facilitate cell-cell interactions [2] and form molecular bridges at synapses to influence synaptic strength and plasticity [3]. CASPR2 is a transmembrane protein with a small intracellular and a large extracellular component (see Figure 1) containing four laminin G domains and two epidermal growth factor-like (EGF-like) domains thought to be involved in receptor-ligand interactions, cell adhesion, migration and differentiation [4, 5]. Unlike most neurexin proteins, the CASPR2 extracellular region also contains a discoidin/neuropilin homology domain and a fibrinogen-like region which mediate cell-cell adhesions and extracellular matrix interactions [6].

**Figure 1.**
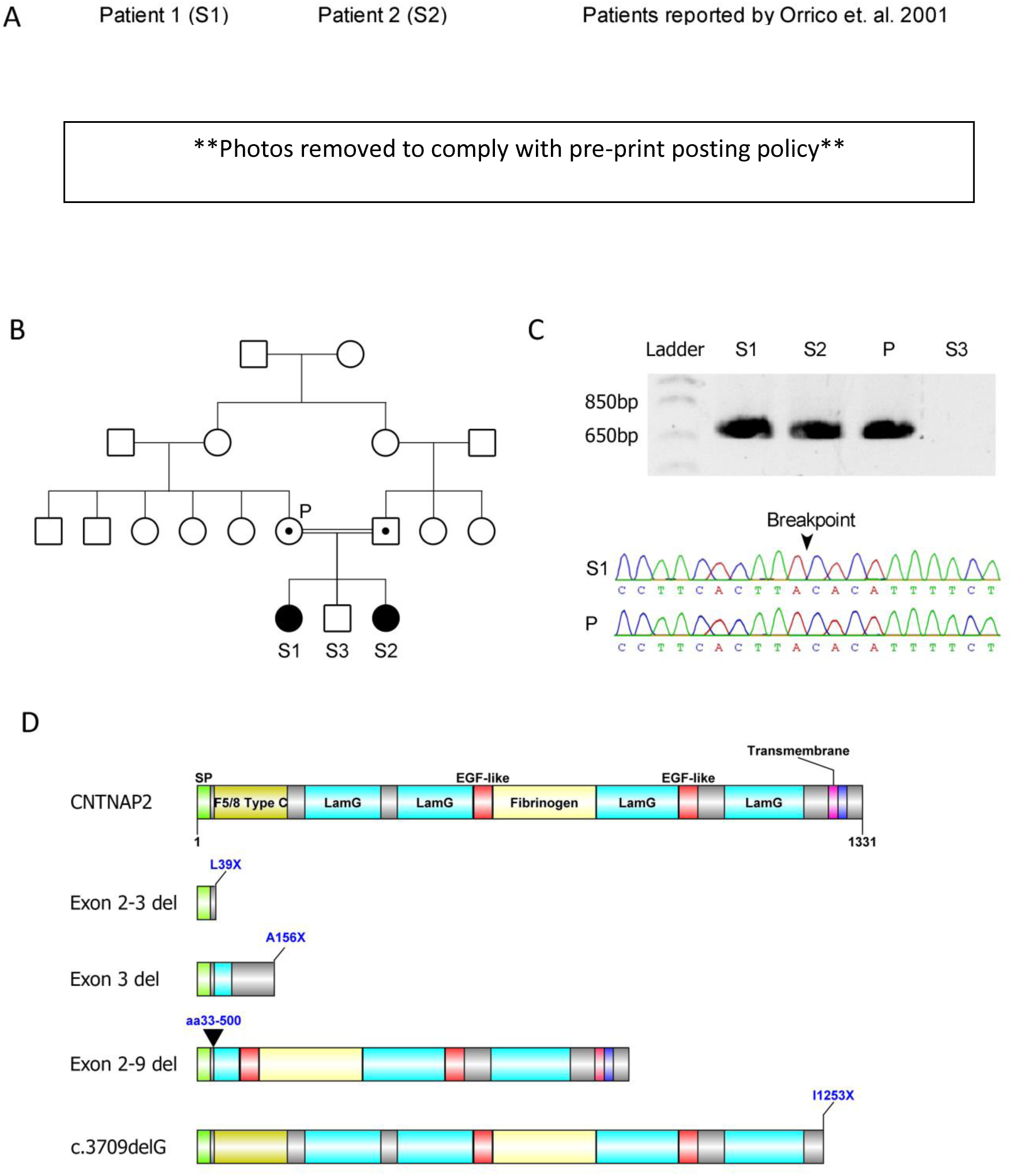
(A) Clinical picture of the affected probands reported in this study. Note dysmorphic facial phenotype, low forehead and bushy eyebrows. Photos of the patients reported in Orrico et. al. are included for comparison [18] (adapted from [18]) (B) Pedigree of the family showing the two affected siblings (filled symbols) with homozygous deletions encompassing exons 2-3 of the *CNTNAP2* gene. The brother is unaffected (unfilled symbol) and has no detected CNV’s. Parents are first cousins and both carry the heterozygous *CNTNAP2* deletion (symbols with dots), but are phenotypically normal. (C) PCR gel of the mutant *CNTNAP2* gene. A ~700 base pair product could be amplified from the homozygous siblings and heterozygous parent, but not from the unaffected sibling. A Sanger sequencing trace shows the location of the deletion breakpoint. (D) The wild type CASPR2 protein and functional consequences of *CNTNAP2* patient mutations on the CASPR2 protein product. CASPR2 is composed of 1331 amino acids with a number of functional domains including a signal peptide (SP), discoidin/neuropilin homology domain (F5/8 Type C), Laminin G domain (LamG), Epidermal Growth Factor like domain (EGF-like), Fibrinogen domain (Fibrinogen), Transmembrane domain (Transmembrane), and Protein 4.1 homologous binding domain (4.1m). The exon 2-3 deletion (reported herein), exon 3 deletion[16] and c3709delG mutation[14] all produce truncated proteins due to early stop codons. The exon 2-9 deletion[13] also produces a truncated protein product, however in this case it is due to an in-frame loss of amino acids 33-500.

*CNTNAP2* is expressed across a number of brain regions throughout development and into adulthood (see [1] for review). In humans, *CNTNAP2* is expressed in the cortex with enrichment in frontal and perisylvian regions, and in the striatum, thalamus and amygdala [7, 8]. This expression pattern recapitulates the cortico-striato-thalamic circuitry known to modulate higher order cognitive processes, including language, reward, and frontal executive function. In mouse models, CASPR2 affects synaptic transmission and dendritic arborisation [9], and both mouse models and human patients have suggested that *CNTNAP2* is important for normal cortical development [10]. Thus it is clear that CASPR2 plays an important role in brain development and function.

## MATERIALS AND METHODS

### Patients

Ethical approval for this study was obtained by the Ethic Committee of the University of Bari, Italy.

### Genetic study

Array-CGH (aCGH) was performed using the BlueGnome ISCA 4x44k Cytochip oligoarray. Copy number variants (CNVs) were identified using the BlueFuse multi v2.3 software (BlueGnome). Human genomic DNA was used as a positive control (Promega #G1471)[11]. A CNV was identified on chromosome 7. To confirm the deletion, DNA was extracted from blood samples of the affected sibs and family controls. PCR primers flanking the deletion were used to amplify DNA and determine the approximate deletion breakpoint. PCR was performed using Taq DNA polymerase kit (Invitrogen) as per supplier's instructions. Following this, a fragment spanning the breakpoint was amplified using the Advantage2 PCR kit (Clonetech, as per supplier's instructions), using the following primers; Fwd: TCCGGTCTTGGCATTGGGAATCATTAGCC, Rev: T CAGCAGAGTGAAGAGACAACCCACAGAGT. This breakpoint spanning PCR yielded a 730bp product from homozygous and heterozygous DNA but not from control samples. This product was then Sanger sequenced to identify the precise breakpoint.

## RESULTS

### Clinical report

Herein, we report a consanguineous family carrying a deletion disrupting the coding region of *CNTNAP2*. Two affected female siblings (Patient 1 and 2) identified in this family were homozygous for the deletion (Figure 1). The affected sisters had one unaffected male sibling and their parents (first degree cousins) were also unaffected. Patient 1 and 2 were born at full term displaying normal height, weight and head circumference with no reported perinatal problems. Since 2 years of age the weight of both patients has been consistently >97^th^ percentile, with Patient 2 classified as severely obese. This has resulted in patient 2 having limited movement (awkward gait), however, neither sibling displays focal motor deficits of the limbs. Additionally, both siblings display a dysmorphic facial phenotype with low forehead and hairline, bushy eyebrows, high arched palate and dental malocclusion (Figure 1A).

Psychomotor development was normal up to 36 months for patient 1 and 18 months for patient 2. At these points, the patients developed trunk swinging motor stereotypies and complex partial seizures. These seizures were characterised by loss of contact, emission of screams, expressions of fright and redness of the face with perilabial cyanosis. Over time the seizures became more frequent and additional features developed including stertorous breathing, loss of consciousness and sometimes generalised tonic-clonic movement or falling to the ground. The seizures were rapidly and completely controlled by Carbamazepine (1000 mg/day) in patient 1. However, for patient 2 Carbamazepine treatment provided only a temporary decrease in the frequency of seizures and she ultimately became unresponsive to it as well as several other anticonvulsants (valproate, phenobarbital, vigabatrine, clobazam, lamotrigine and topiramate). Currently Patient 2 has daily seizures of variable frequency (up to 4–5 episodes per day).

An EEG performed in patient 1 displayed poorly organised background activity without slow and eliptiform abnormalities. In contrast, patient 2 displayed short theta widespread outbursts with pseudo-periodic trend at 20 months of age. During sleep these outbursts were prevalent in the left hemisphere of the medial temporal region. At 4 years and 2 months of age, patient 2 performed another EEG. In this case, background activity was well organized, with outbursts of slow waves (4/5 Hz) sometimes preceded by small amplitude spikes in the right temporal region and spreading to temporal regions of the contralateral hemisphere. During sleep, the anomalies assumed the character of slow waves and slow spike-waves that spread from the right hemisphere to the contralateral side. Recorded seizures showed theta-delta waves and spikes in right temporal hemisphere and vertex, followed by 3 Hz slow activity spread throughout the right hemisphere. At 14 years old patient 2 underwent EEG exams during sleep and showed diffuse epileptiform abnormalities (spikes) in anterior bilateral regions.

An MRI scan of patient 1 did not identify any gross structural abnormalities. However, patient 2 displayed hypoplasia of the cerebellar vermis and hemispheres. Both patients have now been diagnosed with severe intellectual disability according to the Wechsler adult intelligence scale, and have also been classified with autistic spectrum disorder (DSM 5).

Language regression occurred simultaneously with seizure onset in both patients. For patient 1, only syllables were initially lost followed by disappearance of verbal production. Currently, the proband’s language skills are limited to a few isolated words, rarely spoken with communicative intent. Patient 1 also displays vocal tics, reduced relational capacity and autonomy, together with increased stereotypies and clumsiness. Patient 2 had progressive behavioural changes with a slowing in language acquisition, impoverished interpersonal skills and verbal communication, including inertia and apathy. The patient’s mother reported alternating periods of more pronounced symptoms, with periods in which seizures were better controlled and the child appeared to be more attentive and involved, but also more aggressive.

### Genetic analysis

Molecular analysis of the patients was normal for karyotype (46XX) and negative for the presence of sub-telometric deletions. Array-CGH revealed a deletion of at least 127 Kb in chromosome 7q35 (hg18; Chr7:146,082,310-146,189,705). Both affected probands were homozygous for the deletion, while the parents were heterozygous. The unaffected male sibling did not carry this deletion. The exact breakpoints were mapped via PCR and direct Sanger sequencing revealing a 203 kb deletion (hg38; Chr7:146,711,006-146,914,175) spanning exon 2-3 of the *CNTNAP2* gene. *CNTNAP2* expression was assayed in peripheral blood and in transformed lymphoblast cells, but expression levels were too low to reliably detect a signal in patient or control samples (data not shown (see also [12, 13]). Sequence analysis suggested that the loss of exons 2-3 introduces a frameshift into the *CNTNAP2* transcript. This frameshift produces an early stop codon at amino acid 39 (L39X) that would severely truncate the protein product and is likely to lead to nonsense mediated decay (Figure 1C). This suggests that the CNV reported in these patients results in complete loss of functional CASPR2 protein in the homozygous state.

## DISCUSSION

Herein we report two sisters with a homozygous deletion affecting the *CNTNAP2* gene and a phenotype including dysmorphic features, severe mental retardation, early onset seizures, autism spectrum disorder and language impairment. Heterozygous family members carrying this deletion were phenotypically normal. Previously, only three other examples of homozygous loss-of-function *CNTNAP2* mutations have been reported. A search of publicly available exome sequencing data (ExAc database (http://exac.broadinstitute.org/)) did not show any homozygous coding changes in *CNTNAP2*. One SNP in intron 12 (rs113545270) was reported to be homozygous in 76 of 5,077 exomes in the ExAc database. This variant was also reported as a possible loss-of-function change for an alternative transcript variant of CNTNAP2; however there is little evidence that this highly truncated variant is expressed. Furthermore this alternative transcript has been flagged as ‘suspect’ by the ensemble database (http://www.ensembl.org/index.html), suggesting that this is a non-coding change. The first of the reported homozygous *CNTNAP2* deletions consisted of an old-order Amish cohort with a frameshift mutation (c.3709delG) in 18 individuals resulting in the introduction of an early stop codon towards the C-terminus of the CASPR2 protein (I1253X) (Figure 1C)[14]. The protein produced from this mutation retains all the extracellular CASPR2 domains, but is no longer tethered to the neuronal cell membrane and is secreted, suggesting a loss of functional CASPR2 [15]. An unrelated consanguineous family was identified in which three members had a homozygous deletion of exon 3 within the *CNTNAP2* locus [16]. The loss of exon 3 is predicted to introduce an early stop codon (A156X) due to a frameshift mutation. This is likely to result in nonsense mediated decay (NMD) of the RNA product and complete loss of CNTNAP2 protein in these probands (Figure 1C). Finally an in-frame homozygous deletion of *CNTNAP2* exons 2-9 (amino acids 33-500; del.33-500) was reported in two siblings of another unrelated family [17]. Deletion of these amino acids is predicted to result in the partial loss of extracellular domains (Figure 1C). Molecular studies would be needed to determine if this mutant form of CASPR2 retains any functional characteristics, however the loss of almost half the extracellular domain of this protein is suggestive of a severe disruption of function. As in the family reported herein, heterozygous family members were identified in each of these three cases that were phenotypically normal.

In all cases where CASPR2 deficiency was identified, individuals displayed severe intellectual disability and early onset of severe seizures, beginning by 13-36 months of age and often associated with speech/language and behavioural regression (Table 1). Most patients displayed seizures that were refractive to pharmacological or surgical treatments. Pharmacological intervention has only successfully managed seizures in 3 patients (the A156X siblings were responsive to carbamazepine and valproate respectively and patient 1 described herein was responsive to carbamazepine). In all patients, seizures were severe and frequent, with multiple episodes per day being reported in all cases. However, the early seizure onset observed in these patients presents difficulties in distinguishing between phenotypes such as speech/language impairment that could be directly due to *CNTNAP2* mutation or secondary to seizures. For example in I1253X patients, language comprehension was reported to be good before seizure onset around 13-20 months, but all patients showed language regression from 36 months. However, in contrast, in the A156X family the male child did not speak or vocalise, even before seizure onset. Similarly, the del.33-500 siblings only developed a few isolated words. The patients we report herein (L39X) showed language regression occurring simultaneously with seizure onset, until speech became limited to the rare use of isolated words. Taken together, these families suggest that *CNTNAP2* loss results in seizures that may cause language regression, but that it may also be affecting the development of regions of the brain involved in speech and language as impairments in these faculties can occur before seizure onset.

**Table 1.**
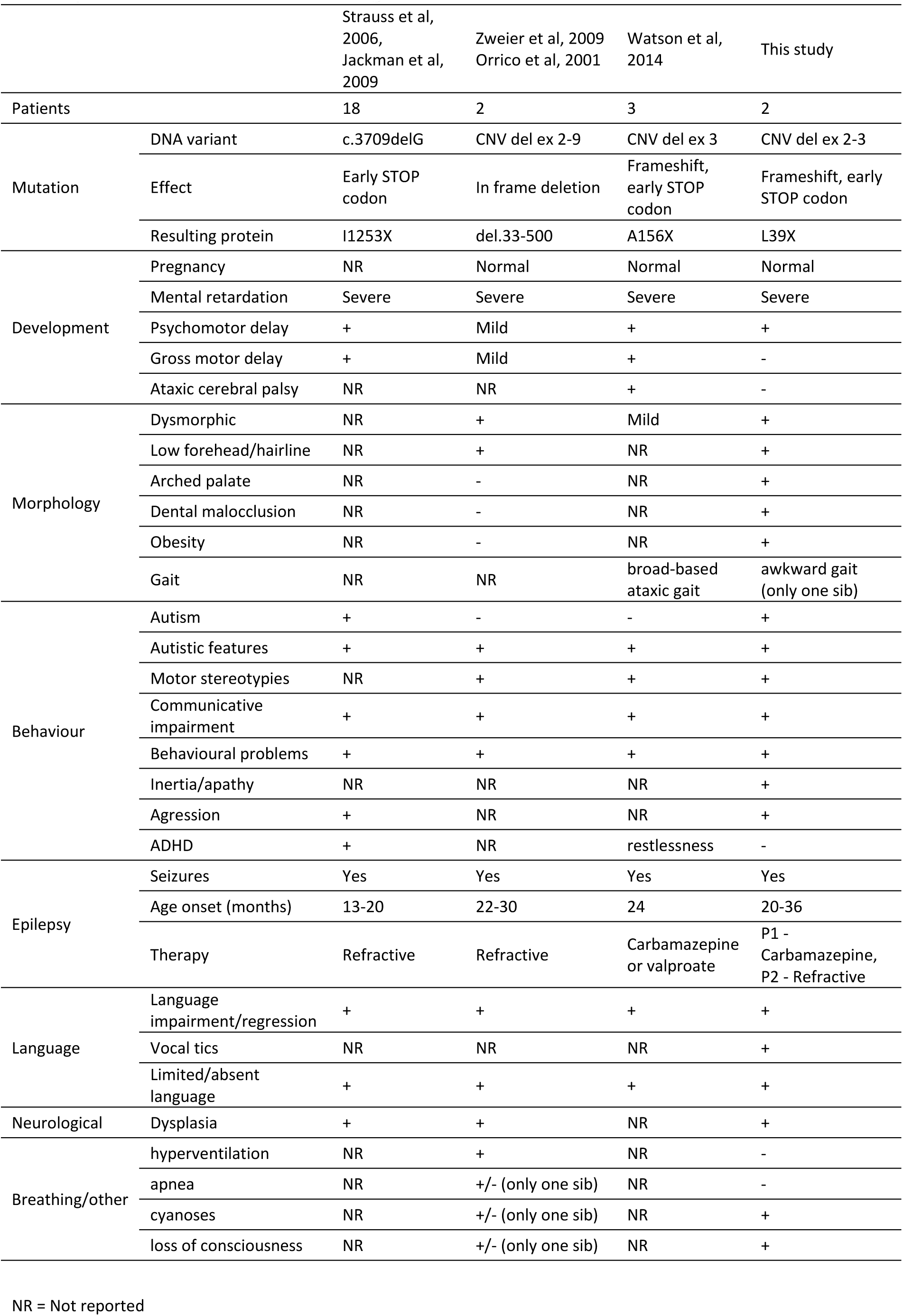
Physical, neurodevelopmental and behavioural features of patients identified with homozygous CNTNAP2 mutations resulting in CASPR2 deficiency disorder.

All patients have also been reported to have some autistic characteristics. Both of the patients described herein are classified as autistic, with impoverished communication skills, communicative impairment, repetitive behaviour, and motor stereotypies. The majority of patients reported with the I1253X mutation were classified as autistic, displaying behavioural deterioration including restricted behaviours, aberrant social interactions and communicative impairment [14]. Zweier et. al. noted that the del.33-500 siblings had limited communicative ability with variable “autistic behaviour and stereotypic movements” [13] and Watson et. al. reported that A156X siblings displayed obsessive compulsive behaviour, stereotypical movements and communicative and speech impairments [16]. Given that *CNTNAP2* remains a top candidate gene with regards to the molecular basis of autism, these homozygous patients would be of great value for studying the links between autism and *CNTNAP2*. Thus there is a need for further in-depth phenotyping of the autistic characteristics of the homozygous patients in these families.

Additional common features between the probands include dysmorphisms, focal cortical dysplasia, behavioural problems and psychomotor delay (Table 1). Dysmorphic features were reported from three of the four mutations (del.33-500, A156X and L39X). Although patient photographs were not available for all families, it was striking to note the facial similarities of the patients described herein and the del.33-500 patients, for whom photos have been published (Figure 1)[18]. Both sets of siblings showed coarse facial features, low forehead, heavy eyebrows and large mouth with thick lips. In future, it will be of interest to determine if this mild dysmorphia is a universal characteristic of CASPR2 deficiency.

Thus while some variability in phenotype remains, it is clear that homozygous loss of *CNTNAP2* results in a syndromic disorder characterised by severe intellectual disability, early-onset drug-resistant epilepsy, reduced or absent language, communicative impairments and autistic phenotypes. To further refine the diagnostic criteria for CASPR2 deficiency disorder (CDD) it will be essential to identify novel patients carrying homozygous *CNTNAP2* mutations by screening cohorts showing core syndromic phenotypes, together with a more detailed phenotypic analysis of existing and novel patients. Furthermore, although the mutations discussed above are predicted to result in a loss of CASPR2 function, cell based and molecular assays will be critical to determine which mutations represent a full vs. partial loss of CNTNAP2 function. This is particularly relevant for mutations such as del.33-500 or I1253X where truncated forms of the protein, retaining some functional domains, are predicted to be expressed. Determining which mutations represent a true loss of function together with increased phenotypic analysis will be of great value in refining the phenotype associated with CASPR2 deficiency disorder, and in understanding the contribution of CNTNAP2 to cognitive development as well as disorders such as epilepsy, autism and language impairment.

## AUTHOR CONTRIBUTIONSs

PRC carried out the molecular genetic studies, NP collected patient samples, TF and AN phenotyped the patients and contributed to the manuscript, PS analysed data and drafted the manuscript and PRC and SCV designed the study, analysed data and drafted the manuscript.

## CONSENT

Written informed consent was obtained from the parent of the patients for publication of this Case report and any accompanying images. A copy of the written consent is available for review by the Editor of this journal

## ACKNOWLEDGMENTS

This work was supported by a Marie Curie Career Integration Grant awarded to S.C.V. and by the Max Planck Society. We are grateful to the family for their consent in undertaking this study.

